# Electrostatics Govern Protein Orientation on Citrate-Capped Gold Nanoparticles

**DOI:** 10.1101/2025.11.07.687172

**Authors:** Suman Tiwari, Akash Kumar Jha, Simran Arora, Debanjana Das, Sri Rama Koti Ainavarapu

## Abstract

Conjugation of proteins with gold nanoparticles (GNPs) is crucial in developing nanoparticle-based drugs, which are used in making targeted delivery systems for therapeutic and diagnostic use. This conjugation is often mediated by covalent gold–thiol (Au-S) bonds involving cysteine residues. However, the influence of other interactions, like electrostatics, on protein-GNP interactions remains poorly understood. While the independent roles of cysteine (thiol interactions) and lysine (electrostatics) are studied, how these two distinct mechanisms couple and mutually influence one another remains unknown. It is particularly unclear how the position and the electrostatics around cysteine residues within the protein structure modulate their interaction with GNPs. Hence, to isolate the electrostatic contribution from Au-S interaction, we systematically investigated the role of positively charged lysine in protein interaction with citrate-capped GNPs using protein L, a cysteine-free antibody-binding protein. Protein L has seven lysine residues, positively charged at pH 7.4. Each lysine was individually mutated to cysteine to create single-cysteine variants. The capping strength of these mutants on citrate-capped GNPs was assessed using various biophysical techniques. Surprisingly, not all cysteine mutants formed stable covalent conjugates. The K7C mutant, in particular, showed strong and irreversible GNP capping, not solely due to the introduced cysteine, but also because of nearby positively charged residues that promoted electrostatic attraction to the negatively charged citrate-capped GNP surface. This critical role of electrostatics was further confirmed using lysine acrylation, which effectively capped lysines and prevented their interaction. Our experimental plan was thus able to successfully decouple the contributions of Au-S bonding from lysine-driven electrostatics. These sequential and cooperative studies underscore that electrostatic and covalent factors are not independent but part of an interconnected process, thereby underscoring the need to consider local charge environments in protein engineering for nanoparticle conjugation.

## Introduction

One of the most exciting applications of nanotechnology is in nanomedicine, where nanoparticles, similar in size to biomolecules, can be designed for cell permeability^1^ and targeted drug delivery.^2^ These applications often utilise various nanoparticle systems, including polymer-based^3,4^ (e.g., polystyrene), lipid-based^5,6^, and inorganic metal-based^7,8^ These applications are being explored for these uses. Among them, gold nanoparticles (GNPs) have attracted significant attention due to their tunable size and shape, enabling targeted drug delivery^9^ and selective accumulation in cells for radiotherapy^10^. These GNPs, ranging from 10 nm to 100 nm, are safely excreted by the kidneys or follow degradation pathways similar to gold salt-based drugs^11^. However, two key challenges remain— maintaining the colloidal stability of GNPs in biological fluids^12,13^, where biomolecular interactions can aggregate GNPs, and achieving efficient accumulation at target sites while ensuring safe excretion or degradation^14,15^. Addressing these challenges requires a deeper understanding of the interface between GNPs and biomolecules such as proteins, lipids, and DNA.

Studying the interaction of GNPs with proteins is particularly important since antibodies, along with cell receptors and transporters, are all proteins^16,17^. Composed of 20 different naturally occurring amino acids, these proteins provide diverse functionalities, which are tuneable to enhance their binding affinity to GNPs and their drug-loading capacity^18^. The protein-GNPs interface is being widely studied to increase the stability of GNPs and to attain a proper orientation of protein on the GNPs surface^19–21^. The most widely studied form of GNP is 10-20 nm citrate-capped GNPs^14^. Towards increasing the stability of GNPs, proteins are sometimes immobilized using a covalent linker that facilitates an Au– linker–protein interaction^22,23^. Linker-based chemistry often causes aggregation of GNPs because the linkers must displace the original protective capping agents to bind to the surface. Since these linkers are typically not effective stabilisers themselves, they fail to prevent the nanoparticles from aggregation, a process that also decreases the loading capacity of the GNPs ^24–26^.

Furthermore, direct immobilisation of proteins through their side chains on GNPs presents its own challenges, where a key question remains: *Is an exposed cysteine sufficient to cap the GNP with protein irreversibly?* ^27^ In a study on the GB3 protein, Fitzkee et al. used NMR to demonstrate that even in the absence of cysteine, GB3 exhibited a strong affinity to the GNPs surface^28^. Further research from the same group revealed that introducing a K13C mutation in GB3 led to a dramatic increase in interaction strength with GNPs^29^. In contrast, a K19C mutation did not show such a dramatic effect, despite both cysteines having similar exposure^30^, which suggests factors other than exposed cysteines drive GB3 interaction with GNPs. A similar study on cytochrome C has shown how cysteine position is important in a protein to immobilise it on a GNP, where one particular double cysteine mutant, K99C-C102 mutant showed an irreversible binding to the protein^31^. Consistent with this, methylation of GB3 lysines drastically reduced its adsorption to citrate-capped GNPs, implicating that lysine residues play a key electrostatic role and are major contributors to the initial binding interface ^32,33^. These studies suggest that additional factors, like exposed lysines beyond cysteine exposure, govern protein immobilisation.

To systematically study protein–GNP interactions, we selected protein L, a bacterial, cysteine null protein that naturally binds antibody light chains, as a model system. While previous studies using proteins such as GB3 and cytochrome C have characterised both electrostatic and Au-S interactions, they generally did not systematically probe the interplay between these factors. Therefore, our work focuses specifically on these potential cross-effects of how local electrostatics around cysteine residues modulate their affinity for gold. To investigate this, our approach is to introduce cysteine mutations at lysine positions with varying solvent exposure. This strategy allows us to test whether exposure alone is sufficient for stable adsorption while simultaneously assessing the effect of altered electrostatics due to lysine replacement and lysine blocking using acrylation. This strategy enables the disentanglement of the contributions of thiol-mediated interaction versus electrostatic interactions, a focus not explicitly addressed in prior work. By immobilising protein L on citrate-capped GNPs, we further created a model interface capable of binding antibodies, demonstrating that surface-immobilised proteins can retain functional recognition. This system thus provides a controlled framework to investigate the principles governing protein adsorption, orientation, and functional retention on nanoparticle surfaces.

## Materials and Methods

### Materials

Chloroauric acid (HAuCl_4_·3H_2_O), dibasic sodium citrate dihydrate ((Na_2_C_6_H_6_O_7_·2H_2_O), 5,5’-dithiobis-(2-nitrobenzoic acid) (DTNB), and 2-mercaptoimidazole (2-MI) were procured from Sigma-Aldrich. Alexa488-NHS-ester dye was procured from Thermo Fisher Scientific. Rabbit IgG Anti-His-Tag Monoclonal Antibody was obtained from Cell Signalling Technology (Danvers, MA, USA).

### Gold nanoparticles synthesis and characterisation

GNPs were synthesised using the citrate reduction method^34^, where 0.5 mL of 25 mM HAuCl_4_ was added slowly to 50 mL of boiling 4.5 mM sodium citrate solution made in Milli-Q water. The solution exhibited a colour change from pale yellow to wine-red, indicating nanoparticle formation. The reaction was maintained at boiling for 10 minutes and then cooled to room temperature under stirring. The synthesised nanoparticles were stored at 4 °C in the dark. The size and shape of GNPs were confirmed using dynamic light scattering (DLS), transmission electron microscopy (TEM), and UV-visible absorption. DLS measurements were performed using a DynaPro NanoStar to determine the hydrodynamic radius of GNPs. Samples were filtered through a 0.45 µM filter before measurement. Data acquisition was carried out at a scattering angle of 90° and analysed using the DYNAMICS software. TEM measurements were performed using a 200 kV Tecnai-20 transmission electron microscope, where a drop of diluted GNPs solution was adsorbed and dried onto a carbon-coated formvar grid (Electron Microscopy Science). Images were analysed to assess particle uniformity and dispersion.

The absorption spectra of GNPs were recorded using MultiSkanGo, with 4.5 mM citrate as the baseline. Measurements were taken over a wavelength range of 300–900 nm to characterise the surface plasmon resonance peak.

### Expression of protein L mutants

Site-directed mutagenesis was performed to construct Cys and Ala mutants of protein L, i.e., protein L K7/23/28/41/42/54/61C and K7/42A in pQE80L (ampicillin-resistant) vector. The plasmids containing the genes encoding for protein L mutants as well as the WT (wild type) variants, with a TEV cleavage site between the N-terminal His_6_-tag and protein, were transformed into *Escherichia coli* BL21 (DE3) cells. Initiating primary cultures from a single colony, 10 mL of LB media was used, grown at 37 °C and 200 rpm for 14 to 16 hours. These cultures were then added to 1 L of LB media with ampicillin, allowing growth at 37 °C until the OD reached 0.4 to 0.6 at 600 nm. Induction of protein expression occurred at 37 °C using isopropyl β-D-1-thiogalactopyranoside (1 mM) for 6 hours. After centrifugation at 6000 rpm, 4 °C for 30 minutes, cell pellets were stored at -80 °C.

Cell pellets were resuspended in PBS buffer (20 mM sodium phosphate, 150 mM NaCl, pH 7.4) with 1X protease inhibitor cocktail (Sigma Aldrich) and lysed by sonication at 4 °C. The resulting supernatant, obtained after centrifugation (17,000 rpm, 45 minutes, 4 °C), was subjected to Ni-NTA chromatography for purification. The resin was successively washed with PBS buffer and 20 mM imidazole (prepared in PBS buffer). Elution of proteins possessing His_6_-tag at the N-terminus was achieved using 250 mM imidazole in PBS buffer. The buffer of the elute was exchanged to 50 mM Tris–HCl, 0.5 mM EDTA, and 1 mM DTT (pH 8.0), and was incubated with 30 µM of TEV protease, at room temperature for 8 hours. Post-cleavage, the reaction mixture was subjected to Ni-NTA chromatography to separate the cleaved His_6_-tag and TEV protease (which was His_6_-tagged) from the target protein.

Purity of the eluted protein was confirmed by SDS-PAGE, where the protein band was observed at the correct size. Final purification involved Fast Performance Liquid Phase Chromatography (FPLC) using Superdex 75 columns (GE Healthcare) on a Bio-Rad Biologic Duo-Flow FPLC system, and this was again verified by SDS-PAGE. The proteins were ultimately stored at 4 °C in PBS buffer.

### Solvent accessible surface area (SASA) calculation

The SASA of the mutated residue was analysed using PyMOL. The wild-type protein structure of protein L was obtained from PDB ID: 1HZ6, and the mutation was introduced at the specified residue position using the Mutagenesis tool of PyMol, selecting the most probable rotamer with minimal steric clashes. To ensure accuracy, the solvent probe radius was set to 1.4 Å (default for water molecules), and calculations were repeated for three independent runs^35^.

### 5,5’-dithiobis-(2-nitrobenzoic acid) (DTNB) study

The DTNB assay was used to quantify free thiol groups in the proteins. Absorbance at 412 nm, corresponding to free 5-thio-2-nitrobenzoic acid formed by the reaction of DTNB with free thiols, was used to determine free cysteine content in proteins. The reaction was performed in 20 mM PBS buffer (pH 7.4) at room temperature. Protein L mutants (20 µM each) were incubated with 0.1 mM DTNB for 30 minutes, and absorbance was measured over a wavelength range of 300–500 nm. Thiol concentration was calculated using a standard curve generated with glutathione (GSH).

### Protein L mutants binding to GNPs

A 10 nM GNP colloidal solution estimated using UV–Vis absorbance and the known molar extinction coefficient of 8.7×10^8^ M^-1^cm^-1^ for 20 nm GNPs,^36^ was centrifuged at 10,000 rpm for 10 minutes and resuspended in 2 µM protein solution (protein L WT/K7C/K42C), then incubated for 12 hours at room temperature. After incubation, free protein was removed from the solution by centrifuging and resuspending the protein-incubated GNPs in citrate solution.

### N-terminal labelling with Alexa dye

For N-terminal labelling of protein L variants (protein L K7/42C) with Alexa488 NHS (N-hydroxy-succinimidyl) ester dye, we prepared a 10 mM stock solution of the dye in DMSO. Protein L variants were incubated with Alexa dye in a 100:1 ratio, respectively, for 1 hour, and free dye was separated from the mixture by using Superdex 75 columns (GE Healthcare) on a Bio-Rad Biologic Duo-Flow FPLC system. The proteins were ultimately stored at 4 °C in PBS buffer in the dark.

### Emission studies

Fluorescence spectra for all samples were recorded using a FluoroMax-3 spectrofluorometer (HORIBA Jobin Yvon). The emission spectra of Alexa 488, conjugated to protein L K7/K42C mutants, were measured in the presence and absence of 10 nM GNPs. For the competitive study, 10 µM of unlabelled protein L WT/K42C/K7C mutants were added to GNPs pre-incubated with labelled proteins (K7/42C) and further incubated for 4 hours. Emission spectra were recorded again after the addition of unlabelled proteins.

Measurement of Alexa 488 fluorescence was done by exciting the samples at 488 nm and recording the emission spectra from 495 to 600 nm. The excitation and emission bandwidths were kept constant within 1 to 2 nm for all samples. For all the measurements, the scan speed was 1 nm/s, and the emission spectrum was averaged for three measurements at identical conditions. All experiments were done at room temperature.

### 2-mercaptoimidizaole (2-MI) assay

2-MI is an organo-thiol compound that induces GNP aggregation by removing the reversible capping agents from the GNP surface. A 2 mM stock solution of 2-MI was prepared in water, and a final concentration of 60 µM was added to both bare GNPs and GNPs pre-incubated with protein L WT/K7C/K42C. The addition of 2-MI was done at time intervals of 0, 4, 8, 16, and 24 hours of incubation. Before and instantly after each addition of 2-MI, the absorption spectra of GNPs were recorded in the range of 300–900 nm.

### Isothermal Calorimetry (ITC)

All ITC experiments were conducted at 25 °C (298 K) using a MicroCal PEAQ-ITC instrument (Malvern Panalytical) located at the Department of Biosciences and Bioengineering (BSBE), IIT Bombay, Mumbai. Before titration, both protein and nanoparticle samples were prepared in ultrapure water to ensure buffer consistency and eliminate background heat effects. Diluted solutions of protein L and its variants (WT, K7C, K42C, K7A, and K42A) were used as titrants, along with variable concentrations (40 µM) of IgG antibody (Ab) for comparative analysis. The specific molar ratios of protein-to-nanoparticle and protein-to-antibody used in each titration were optimised to achieve saturation in calorimetric signal by the end of the experiment, as detailed in Table 2. To account for non-specific heat changes, control titrations (blank runs) were performed by injecting nanoparticle solutions into ultrapure water, and the resulting heat of dilution was subtracted from experimental data. During each titration cycle, 3 µL of the protein solution was injected into the sample cell over a period of 6 seconds, followed by an equilibration interval of 150 seconds to allow thermal stabilisation.

The resulting thermograms were baseline-corrected using the control data, and the integrated heat values were fitted to a “one set of sites” binding model, which assumes independent and identical binding sites. Data analysis was carried out using MicroCal PEAQ-ITC Analysis Software (version 1.41), yielding thermodynamic parameters including the binding stoichiometry (N), equilibrium dissociation constant (K_d_), enthalpy change (ΔH), and entropy change (ΔS). These measurements provided quantitative insights into the binding affinity and thermodynamic characteristics governing the interactions between protein variants and nanoparticles or antibody targets.

### Acrylation of lysines

Lysine residues were acylated using N-hydroxysuccinimide (NHS)-acrylate. The protein was incubated with a 10-fold molar excess of NHS-acrylate for 2 hours at pH 8.0 in PBS buffer. Following the reaction, the acrylated protein was separated from unreacted NHS-acrylate by size-exclusion chromatography using a Superdex 75 column (GE Healthcare) on a Bio-Rad Biologic Duo-Flow FPLC system. Successful protein acrylation was confirmed using Electrospray Ionisation Mass Spectrometry (ESI-MS).

## Results

### Characterisation of GNPs

TEM images (Figure 1a) showed that the GNPs were uniformly spherical. The size distribution analysis from TEM (Figure 1c) indicated an average radius of 10 ± 2 nm, whereas DLS measurements gave a larger hydrodynamic radius of 15 ± 3 nm. The higher value observed in DLS arises from the solvation layer and possible inter-particle interactions in solution, which contribute to the apparent hydrodynamic size. A distinct surface plasmon resonance (SPR) peak at 520 nm in the UV–visible absorption spectrum (300–900 nm) further supported the uniformity and stability of the nanoparticles (Figure 1b).

**Figure 1.**
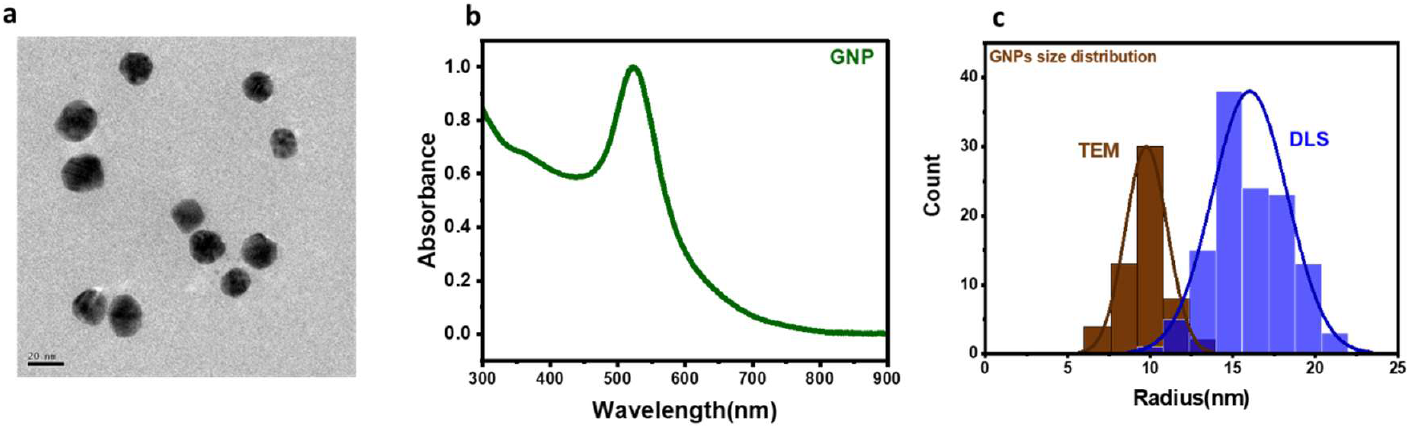
Characterisation of gold nanoparticles (GNPs). (a) TEM image showing uniformly spherical GNPs. Scale bar: 20 nm. (b) UV–visible absorption spectrum of GNPs showing a sharp surface plasmon resonance peak at 520 nm, characteristic of ∼20 nm gold nanoparticles. (c) Size distribution histograms obtained from TEM (brown) and DLS (blue). TEM analysis gave an average radius of 10 ± 2 nm, while DLS showed a larger hydrodynamic radius of 15 ± 3 nm due to the solvation layer surrounding the particles in aqueous medium.

### Free thiol estimation in protein L mutants and SASA values

SASA (solvent accessible surface area) values of cysteine residues are summarised in Table 1. To experimentally assess thiol accessibility, a DTNB (5,5’-dithiobis-(2-nitrobenzoic acid)) assay was performed (Figure 2. b). DTNB, also known as Ellman’s reagent, reacts with free thiol (-SH) groups to produce a yellow-coloured 5-thio-2-nitrobenzoate (TNB) anion, which can be quantified by measuring absorbance at 412 nm. The intensity of the colour change or absorbance at 412 nm is directly proportional to the concentration of free thiols in the sample^37^. The absorbance values at 412 nm for 20 µM protein L K7C and K42C mutants were 0.46 and 0.38, respectively, indicating the presence of accessible free thiol groups. These values were comparable to the absorbance of 20 µM reduced GSH (glutathione) at 0.5. In contrast, all other mutants, including K23C, exhibited negligible absorbance, despite their relatively higher SASA (Figure 2.c). Based on the combined analysis of SASA values and DTNB assay results, protein L K7C and K42C were selected as the cysteine mutants for further binding studies with GNPs, with protein L WT serving as the control.

**Table 1.** Solvent accessible surface area (SASA) values of the cysteine residue of protein L mutants as calculated using PyMol. Protein L K7C and K42C exhibited the highest surface exposure.

| Protein | SASA(Å<sup>2</sup>) |
| --- | --- |
| K7C | 125.1 |
| K23C | 100.5 |
| K28C | 63.5 |
| K41C | 69.5 |
| K42C | 185.3 |
| K54C | 70.5 |
| K61C | 75.5 |

**Table 2.** Thermodynamic parameters of protein L WT, K7/42C interaction with GNPs, as obtained from the ITC titration.

|  | PLWT | K7C | K42C |
| --- | --- | --- | --- |
| $K_d(M)$ | $514 \times 10^{-9} \pm 50 \times 10^{-9}$ | $487 \times 10^{-9} \pm 80 \times 10^{-9}$ | $1010 \times 10^{-9} \pm 95 \times 10^{-9}$ |
| $\Delta H(kcal/mol)$ | $-588 \pm 34$ | $-312 \pm 29$ | $-518 \pm 55$ |
| $\Delta G(kcal/mol)$ | -8.58 | -8.61 | -8.18 |
| $-T\Delta S(kcal/mol)$ | 579 | 303 | 509 |
| N-site | $110 \pm 5$ | $113 \pm 8$ | $98 \pm 9$ |

**Figure 2.**
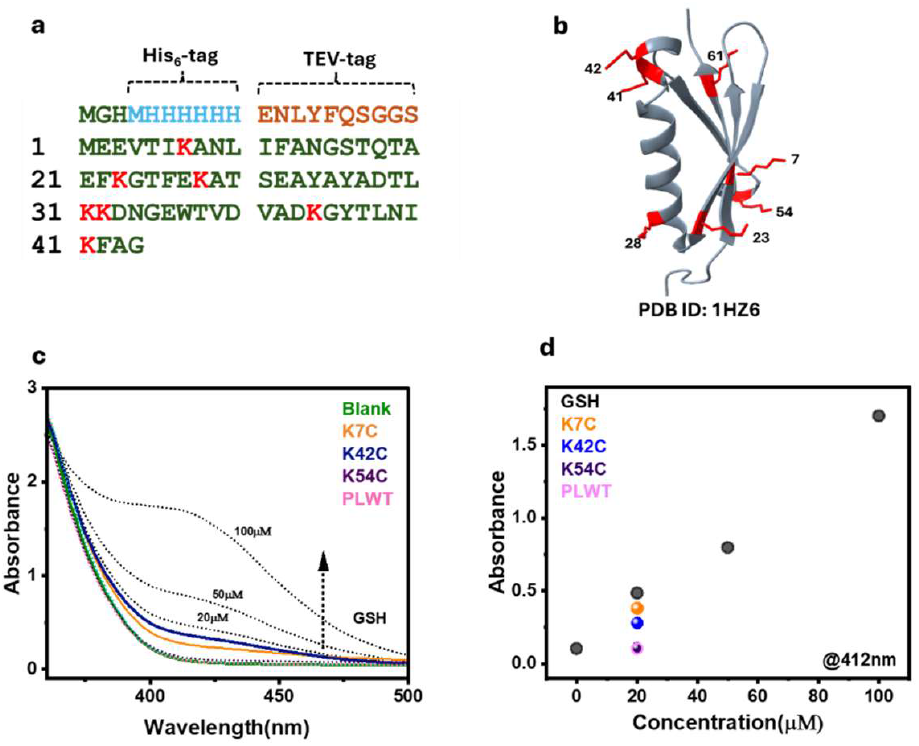
Estimating the surface accessibility of mutated cysteines of protein L. **a**. Amino acid sequence of protein L WT, with lysines highlighted in red, and His_6_ tag and TEV tag of the construct highlighted in blue and orange, respectively. **b**. Crystal structure of protein L (PDB ID: 1HZ6), with lysines highlighted in red and their residue numbers labelled next to them. **c**. UV-Visible absorption spectra of DTNB as a blank, with different concentrations of GSH represented by black dotted lines, and with different mutants of protein L (K7/28/42/54C), along with the wild-type protein L (PLWT), shown as coloured solid lines. **d**. Absorbance at 412 nm vs. concentration plot, where a standard curve of TNB absorbance at different concentrations of GSH is shown in grey. The absorbance values of 20 µM protein L and its mutants are shown. The protein L mutants K7C and K42C exhibited absorbance values of 0.46 and 0.38, respectively, indicating significant exposure.

Among the cysteine mutants, K7C and K42C exhibited the highest surface exposure, with SASA values of 125.1 Å^2^ and 185.3 Å^2^, respectively, as determined using PyMol^35^. The K23C mutant displayed a SASA value of 100 Å^2^.

### Surface energy transfer assay

Surface energy transfer (SET) occurs when an excited fluorophore transfers its energy non-radiatively to the conductive surface of a GNP, leading to fluorescence quenching. This process is driven by the strong dipole-surface interaction between the fluorophore and the free electron oscillations (surface plasmons) of the GNP, which efficiently dissipate the energy. Unlike Förster resonance energy transfer (FRET), which follows a d^−6^ dependence, SET exhibits a d^−4^ distance dependence due to its dipole-surface coupling mechanism^38,39^. Here, we utilise SET of Alexa 488-labelled proteins as a fluorescence-based monitor to determine whether the proteins can be displaced from the GNP surface, where fluorescence recovery indicates protein replacement. N-terminal Alexa 488 labelled K7C/K42C protein L mutants, respectively, showed quenching of fluorescence when incubated with GNPs (Figure 3. c & d). After 4 hr incubation of labelled protein L K42C pre-incubated with GNPs with 10 µM unlabelled protein L WT/K7C/K42C, the fluorescence of labelled K42C mutant recovered by 50%. On similar incubation of labelled Protein L K7C pre-incubated with GNPs, the fluorescence was not recovered at all. The same is represented as a bar graph in Figure 3. e. This hinted at a non-replaceable irreversible conjugation of protein L K7C around GNPs.

**Figure 3.**
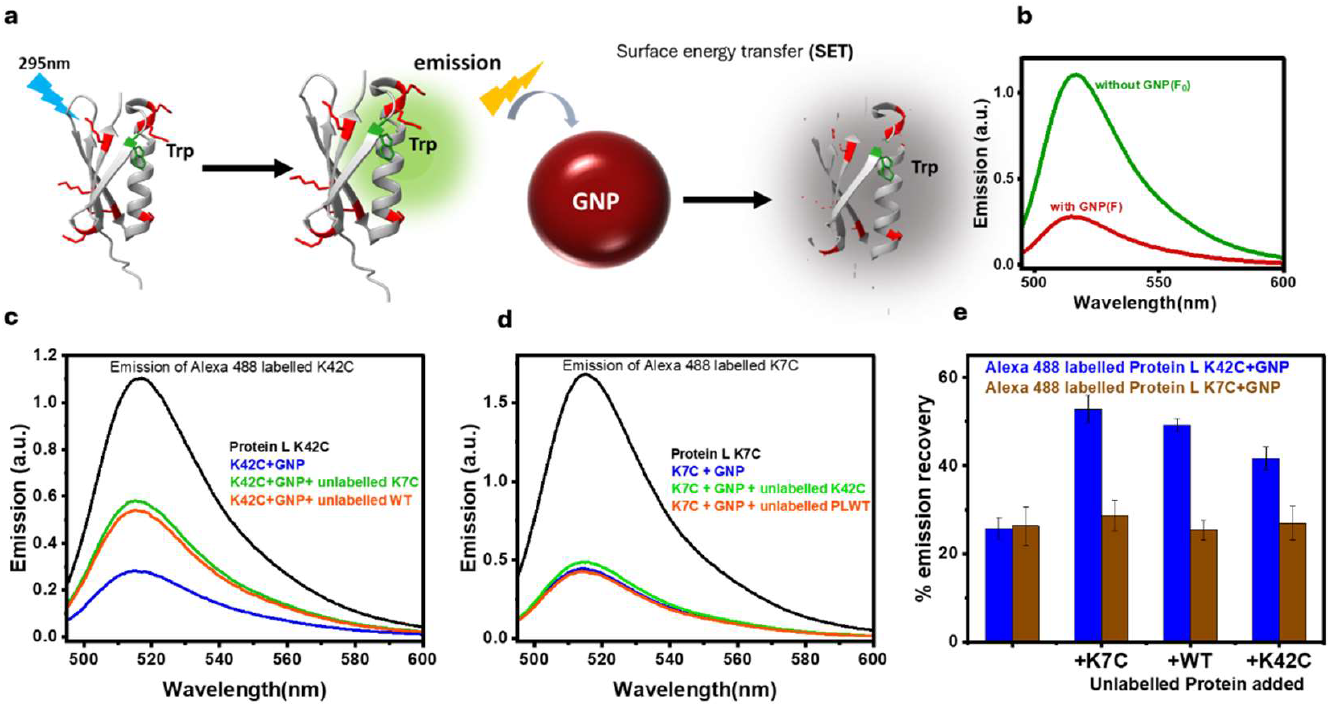
Quenching of tryptophan and Alexa 488-labelled fluorophore of protein L mutants upon equilibrating with GNPs. **a**. Schematic representation of surface energy transfer (SET) from excited fluorophore in the presence of GNPs. **b**. Emission of Trp in the presence and absence of GNPs, showing a significant reduction in emission with GNPs. **c**,**d**. Emission spectra of Alexa 488-labelled at the N-terminus of protein L K42C and K7C in the presence and absence of GNPs. The emission after the addition of other proteins to the GNP-bound protein is shown in green and orange **e**. Percentage of emission recovery vs. protein added for protein L K42C with GNPs (brown) and protein L K7C with GNPs (blue), demonstrating significant emission recovery for protein L K42C.

### 2-MI (2-Mercapto Imidazole) assay

2-MI induces rapid aggregation of GNPs, leading to notable changes in their optical properties. A colloidal solution of 20 nm sized GNPs exhibits a distinct SPR absorption peak at 520 nm. Upon aggregation triggered by 2-MI, the SPR peak broadens, accompanied by a visible colour change from wine red to dark blue. However, when GNPs are irreversibly capped by a protein or another capping agent, 2-MI does not cause immediate aggregation (Figure 4. a,b).

**Figure 4.**
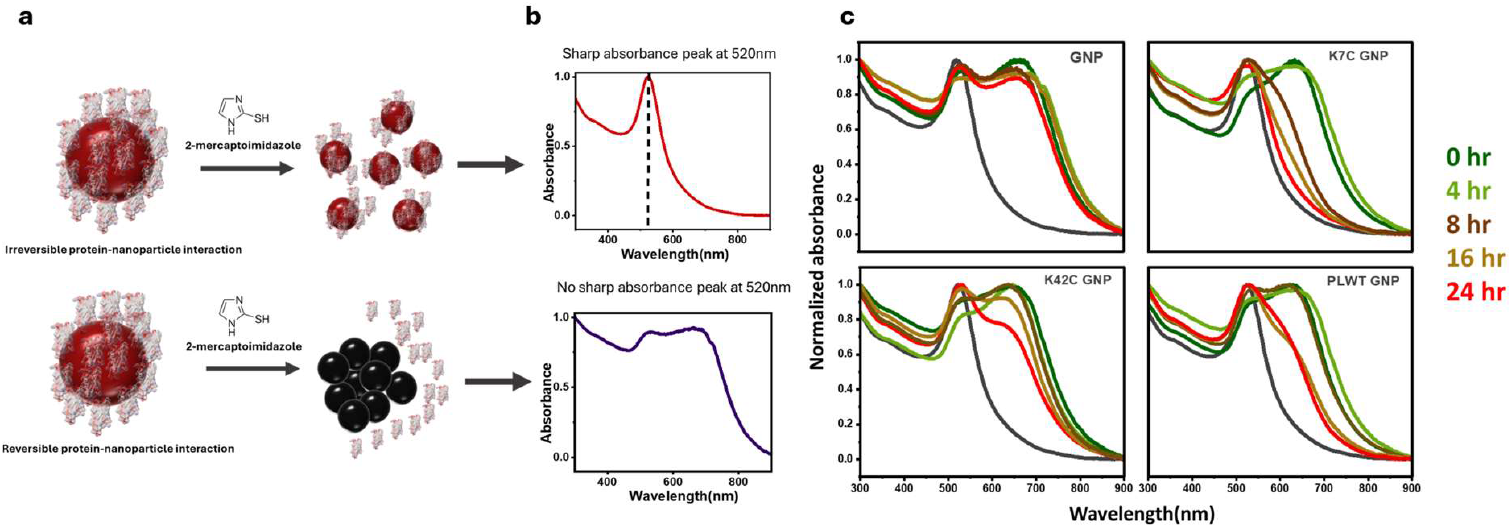
Assessing the irreversible capping of the GNPs with protein L mutants. **a**. Schematic representation of the interaction between 2-mercaptoimidazole (2-MI) and GNPs in the case of irreversible capping, where GNPs remain colloidal, and in the case of reversible capping, where GNPs aggregate. **b**. UV-visible absorption spectra of GNPs in the presence of 2-MI, showing surface plasmon resonance (SPR) at 520 nm in the case of irreversible capping, while in the case of reversible capping or aggregation, the spectrum broadens. **c**. UV-Visible absorption spectra of GNPs, protein L K7C/K42C, and WT with GNPs after the addition of 2-MI at multiple time intervals (0, 4, 8, 16, and 24 hours), shown as green to red-coloured solid lines. protein L K7C with GNPs shows no aggregation after 16 hours.

GNPs incubated with wild-type protein L (WT) for up to 8 hours exhibited complete and instantaneous aggregation upon the addition of 2-MI. However, at longer incubation times of 16 and 24 hours, the extent of aggregation was significantly reduced. In contrast, GNPs incubated with protein L K42C at any time point between 0 and 24 hours displayed immediate and substantial aggregation upon 2-MI addition. Interestingly, after 4 hours of incubation with protein L K7C, a notable reduction in aggregation was observed, and after 24 hours, 2-MI no longer induced aggregation (Figure 4. c).

These findings suggest that protein L K7C forms a relatively stronger and more irreversible capping interaction with GNPs compared to protein L WT. Additionally, protein L K42C exhibited the weakest interaction with GNPs.

### Thermodynamics of protein WT/K7C/K42C interaction with GNPs

To investigate the thermodynamics of protein binding to GNPs, ITC measurements were conducted for the three protein variants, protein L WT, K7C, and K42C. The dissociation constant (K_d_), enthalpy change (ΔH), Gibbs free energy (ΔG), entropy contribution (-TΔS), and binding stoichiometry (N) are summarised in Table 2. The thermodynamic curve of differential power (DP) versus time (min) indicates saturation of thermal changes when protein in the injection cell is mixed with GNPs in the sample cell (Figure 5). All three proteins exhibited a similar thermodynamic curve, exhibiting thermal saturation at nearly the same time.

**Figure 5.**
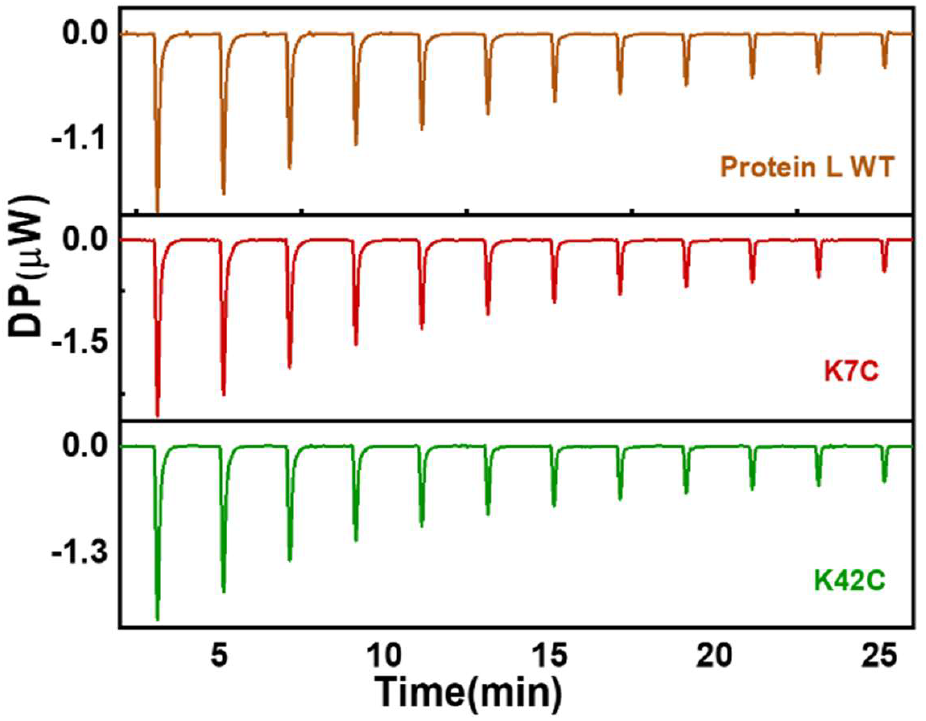
ITC experiments probing the binding energetics of protein L mutants onto GNPs. Corrected differential power (DP) vs. time plot of protein L WT (brown), K7C (red), and K42C (green) titrated against GNPs. The thermodynamic saturation time points for the three titrations are similar, with nearly identical exothermic responses, as indicated by the DP values.

The binding affinity, as indicated by the K_d_ values, varied across the three proteins. Protein L WT exhibited the strongest binding affinity (K_d_ = 487 × 10^-9^ M), which was similar to the binding affinity of Protein L K7C (K_d_ = 514 × 10^-9^ M). In contrast, protein L K42C displayed a significantly weaker binding affinity (K_d_ = 1010 × 10^-9^ M), suggesting a lower stability of the complex.

Binding stoichiometry (N) values indicated that approximately 110–113 protein molecules bind per nanoparticle. Overall, these ITC results suggest that protein L K7C and WT have similar but higher binding affinities, while protein L K42C binds with the weakest affinity.

The alanine mutants of the protein L i.e., K7A and K42A, were made to understand the contribution of Cys in binding affinities. In thermodynamic investigations of the mutants, K7A showed a stronger affinity with a K_d_ value of 140×10^-9^ ± 62×10^-9^ M, and K42A showed a weaker affinity with K_d_ value of 265×10^-9^ ± 57×10^-9^ M (SI Figure 3, Table 2). Although WT showed weaker binding than both K7A and K42A, the relative trend was consistent, with K42 mutations reducing affinity compared to K7. These results highlight residue-specific contributions, but the higher affinities of the alanine mutants likely arise from non-native stabilisation at the nanoparticle interface. Thus, the mutants are useful mechanistic probes, whereas the WT remains the native binding standard for binding.

To test whether alanine mutants bind weaker than PLWT, we used a 2-MI assay. After 24 hours with GNPs, the alanine mutants showed higher aggregation than PLWT when 2-MI was added (SI Figure 4). To verify lysine contribution, we blocked lysines with NHS-acrylate. In the follow-up 2-MI study, acrylated K7C and K42C showed similar aggregation in the presence of 2-MI, unlike the non-acrylated state, where K7C showed no aggregation. Together, the data indicate lysine-dependent stabilisation of GNP capping under these assay conditions (SI Figure 5, 6; SI.1, SI.2).

### Binding affinity of GNP-immobilised K7C to Ab

Protein L is named for its ability to bind the κ light chain of antibodies. The thermodynamic investigation(Table 3) of Rabbit-IgG-Anti-His antibody (Ab) binding to the κ1 light chain with monomeric, free protein L K7C yielded a K_d_ of 1.10 × 10^-6^ M, consistent with reported single-domain affinities in the sub-micromolar range under solution conditions (≈1.3–1.6 × 10−^6^ M at pH 8.0)^40^. In DP traces, Ab binding to bare gold nanoparticles saturated by ∼20 min with DP values of 0–12 µW, indicative of nonspecific adsorption, whereas protein L K7C with Ab and GNP-immobilised protein L K7C with Ab showed reproducible saturation with lower DP values (0–6 µW) (Figure 6). Notably, immobilisation of monomeric protein L K7C on GNPs showed apparent enhancement relative to the free K7C in this assay (K_d_ = 0.18 × 10^-6^ M). However, literature values for multi-domain (native) protein L ^40,41^ measured in immobilised forms are nanomolar (K_d_ ∼1–7 × 10^-9^ M for IgGκ/κ-chain), much tighter than either free or GNP-immobilised single-domain measurements.

**Figure 6.**
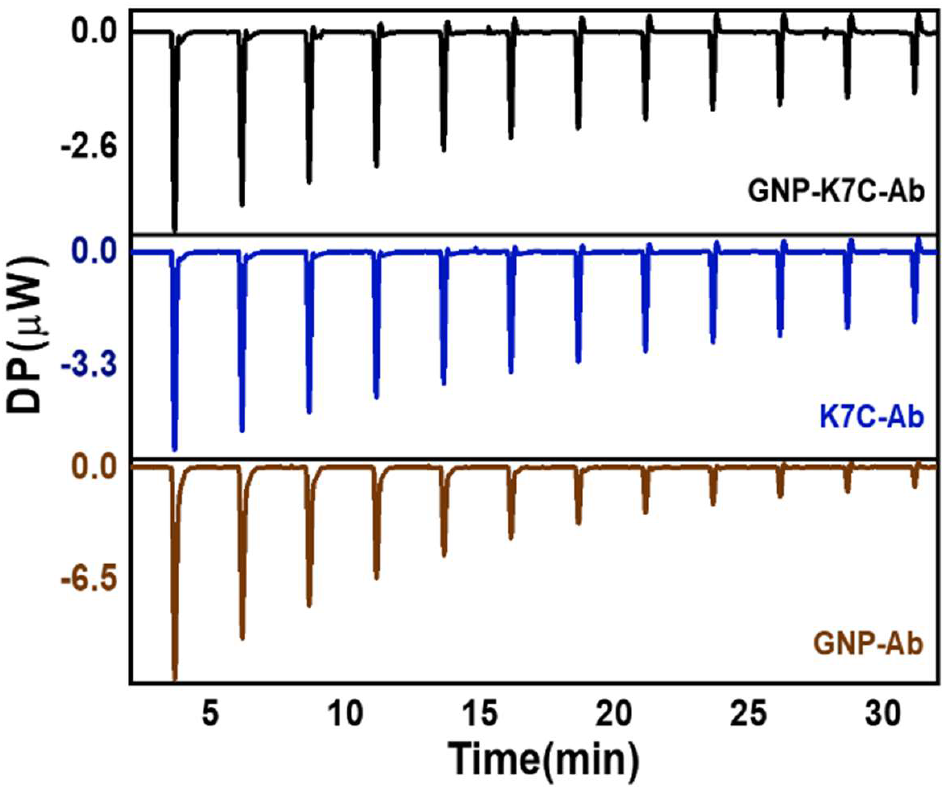
ITC experiments probing the binding energetics of Ab onto GNPs capped with protein L K7C. Corrected differential power (DP) vs time plot of GNP coated with protein L K7C titrated against Antibody (Ab) (in black), and K7C titrated against Ab (blue), and citrate capped GNPs titrated against Ab (brown). The thermodynamic saturation time point of GNPs with Ab is fastest with the highest DP value change, whereas GNPs coated with protein L K7C and only protein L K7C titrated against Ab show nearly similar saturation time points and DP values.

**Table 3.** Thermodynamic parameters of protein L WT, K7/42C interaction with GNPs, as obtained from the ITC titration.

|  | GNP K7C-Ab | K7C-Ab | GNP-Ab |
| --- | --- | --- | --- |
| **$K_d$(M)** | $0.18 \times 10^{-6} \pm 71 \times 10^{-9}$ | $1.10 \times 10^{-6} \pm 96 \times 10^{-9}$ | $0.25 \times 10^{-6} \pm 67 \times 10^{-9}$ |
| **$\Delta H$(kcal/mol)** | $-608 \pm 81$ | $-1500 \pm 106$ | $-2770 \pm 318$ |
| **$\Delta G$(kcal/mol)** | -38.5 | -34.1 | -37.7 |
| **$-T\Delta S$(kcal/mol)** | 570 | 1470 | 2730 |
| **N-site** | 4 | 6 | $10 \pm 2$ |

These findings indicate that cysteine-mediated immobilisation on GNPs substantially enhances the binding capacity of monomeric protein L relative to its free form, most likely by improving orientation and accessibility at the nanoparticle surface. Although the affinity approaches values reported for multidomain protein L, it remains lower, consistent with the absence of oligomeric cooperativity and multivalent avidity that stabilise tetramer–antibody interactions.

## Discussion

This study provides insights into the interaction of proteins with the surface of GNPs. The 20 nm citrate-capped GNPs, characterised by DLS, TEM, and UV-visible absorption studies, were chosen as a model system. Protein L, which lacks native cysteine residues, was selected, and lysine residues were individually mutated to cysteine. Among the seven lysine-to-cysteine mutants, SASA calculations and DTNB assays identified protein L K7C and K42C as possessing accessible thiol groups, making them ideal candidates for protein-GNP interaction studies.

The SET assay provided the first insight into protein-GNP interaction stability. Protein L K7C exhibited a strong, non-replaceable interaction, as indicated by the lack of fluorescence recovery, whereas protein L K42C displayed a weaker, more dynamic interaction. These observations were further validated by the 2-MI assay, which demonstrated that protein L K7C formed an irreversible capping layer, preventing GNP aggregation over time. In contrast, protein L K42C allowed immediate aggregation. Despite both mutants having similar thiol exposure, only K7C formed an irreversible coating, suggesting that cysteine exposure alone is insufficient for strong GNP capping.

ITC further confirmed the interaction strength. Protein L WT and K7C exhibited similar binding affinities, both significantly stronger than K42C. The comparable interaction of WT and K7C during ITC experiments can be attributed to the experimental time frame. The 2-MI assay indicated that cysteine-mediated binding strengthens beyond six hours, suggesting that short-term ITC measurements primarily reflect electrostatic interactions. This observation was further confirmed by comparing the binding affinities of the alanine mutants, i.e., K7/42A, where K42A had a weaker affinity towards GNPs than K7A.

The combined results from SET, 2-MI, and ITC suggest that while protein L WT and K7C initially exhibit similar interaction strengths, K7C eventually achieves irreversible capping, whereas K42C fails to match the capping strength of WT. This difference is attributed to the presence or absence of the most exposed lysine residue (K42). The significantly weaker affinity of both K42C and K42A mutants further confirms the critical role of K42, a finding consistent with established models where K-to-A mutagenesis of exposed lysines abrogates electrostatic binding to GNPs^29^. Since protein L is overall negatively charged (-6), its interaction with negatively charged citrate-capped GNPs is driven by localised positive charges on lysines. The SASA analysis indicated that K42 is the most exposed lysine (SI Table 1), playing a crucial role in mediating the initial electrostatic interaction^32,33^ between the protein and GNPs.

Our findings are consistent with a dual-interaction model reported in the literature, where protein adsorption is not exclusively driven by cysteine-mediated chemisorption^30^. Studies on proteins like cytochrome c show that cysteine-driven binding depends on the residue’s position and solvent exposure^31^. Our 2-MI assay for K7C and K42C aligns well with this finding. In our assay, the difference in binding reversibility between the two proteins established the critical role of electrostatic docking. This result further indicated that the specific cysteine-mediated interaction is a slow process, which strengthens over hours rather than acting as an immediate driver. Reports on proteins, such as lysozyme^42^ and Human Serum Albumin (HSA)^43^, demonstrate that adsorption is mediated by “positive patches” of lysine and arginine residues. This principle was further proven using a cysteine-null variant of GB3, where chemical methylation of its lysines completely inhibited protein attachment to GNPs^32,33^. This body of work directly supports our data. We found that the mutation of the exposed most lysine in protein L (K42A/K42C) resulted in a significant loss of affinity. This observation confirms that K42 acts as the primary electrostatic docking site for protein L.

Our study provides a systematic investigation into why a certain location of a cysteine residue in a protein impacts its stable capping around GNPs. We separated lysine-mediated adhesion from position-specific cysteine interaction within one protein scaffold by using lysine acrylation and modifying lysine to cysteine/alanine mutations. This approach explained two key points: first, that the initial line of interaction between proteins and GNPs is driven by the most positively charged surface (lysine-rich region) of the protein, which is a reversible interaction. Second, cysteine-mediated interaction is achieved at a later stage after the initial electrostatic-mediated interaction and is an irreversible interaction, forming a stable protein coating.

One natural extension of this idea has been the attempt to directly conjugate antibodies to GNPs. Antibodies are particularly attractive because, in addition to forming stable protein–nanoparticle complexes, they inherently carry biological specificity through their Fab domains. This property allows antibody-coated GNPs to selectively recognise and accumulate in target cells, providing opportunities for precision delivery of therapeutic cargo or enhancing local radiation effects^44^. Additionally, Driskell *et*.*al*. have immobilised anti-HRP IgG antibodies directly on GNPs and shown that antibodies with thiolated lysines showed irreversible binding with GNP and also showed functionality as they bound to HRP. This approach was performed to show if antibodies can be immobilised on GNPs with tunable orientation and maintain their functionality^45–47^. However, other reports caution that lysine thiolation can also modify residues within or near the antigen-binding region, which may compromise antibody activity or alter orientation on the nanoparticle surface^48^. Thus, while lysine thiolation is an effective strategy for achieving strong and stable attachment, careful optimisation is necessary to balance immobilisation efficiency with preservation of antibody function.

In this context, protein L, which binds to the light chain of antibodies, offers a useful model for examining whether immobilisation on GNPs preserves antibody-binding activity. To assess whether protein L K7C, once immobilised on GNPs, retains its ability to bind antibodies—a key function of protein L—we performed thermodynamic binding studies. ITC revealed that monomeric protein L K7C immobilised on GNPs retained its ability to bind antibodies (K_d_ = 0.18 × 10^−6^ M). The thermodynamic analysis confirmed that protein L K7C forms a highly stable complex when immobilised on GNPs, preserving its biological function.

Thus, by systematically decoupling the electrostatic (lysine) and thiol-mediated (cysteine) contributions, this study demonstrates that these two mechanisms are not independent but are, in fact, strongly coupled factors in protein-GNP immobilisation. We establish that the electrostatic environment is a critical prerequisite that modulates the efficacy of subsequent thiol capping, an understanding that offers key insights for designing more stable protein-GNP conjugates.

## Conclusion

Our findings demonstrate that protein L K7C exhibits the strongest and most irreversible capping on GNPs, attributed to two key factors: (i) the retention of its most exposed lysine (positively charged), ensuring a strong electrostatic interaction, and (ii) the presence of an accessible cysteine, which enables irreversible binding. In contrast, protein L K42C, despite having an exposed cysteine, lacked the necessary lysine-driven electrostatic interaction and failed to achieve stable GNP capping. Protein L WT, while capable of forming a strong electrostatic interaction, lacked the thiol-mediated irreversible attachment observed in K7C.

Furthermore, the ability of irreversibly capped protein L K7C to bind antibodies highlights its potential for biomedical applications. This property could be leveraged for antibody-functionalized GNPs targeting specific receptors on cancer cells, enabling selective GNP accumulation for radiotherapy^49,50^.

Overall, this study elucidates the coupled roles of lysine and cysteine in protein immobilisation on GNP surfaces. It demonstrates that electrostatic (lysine) and thiol (cysteine) effects are not independent factors but are strongly interconnected, a phenomenon that has lacked systematic characterisation. We show that the initial electrostatic attraction is a critical prerequisite that governs the efficacy of the final thiol-mediated covalent bonding. An understanding of this coupling provides insights for designing protein-GNP conjugates with enhanced stability and biomedical applicability.

## Acknowledgments

S.R.K.A. acknowledges financial support from the Department of Atomic Energy (DAE), India (project no. 12-R&DTFR-5.10-0100). The authors thank Prof. Ashutosh Kumar (IIT Bombay) for providing access to the Isothermal Titration Calorimetry (ITC) instrument.

## Notes

### Competing Interest Statement

The authors have declared no competing interest.

