## supplementary figures tables for "Electrostatics Govern Protein Orientation on Citrate-Capped Gold Nanoparticles"

**Author ORCIDs**

Suman Tiwari: 0009-0003-2913-2947

Akash Kumar Jha: 0000-0002-4580-1480

Simran Arora: 0000-0002-6423-4442

Debanjana Das: 0000-0002-3178-7526

Sri Rama Koti Ainavarapu: 0000-0002-1646-2731

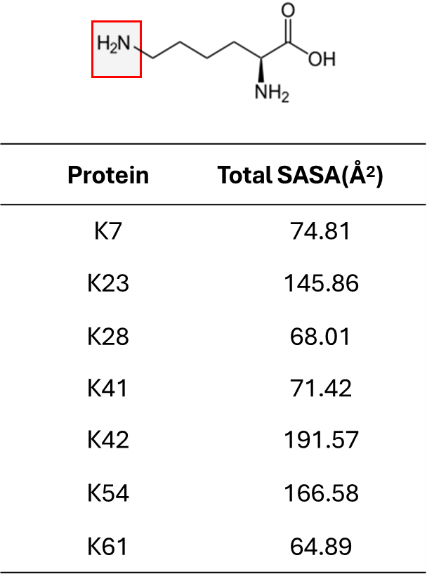

**SI Table 1.** Solvent accessible surface area (SASA) of lysine residues in protein L. The table lists the calculated total SASA values (in Å²) for individual lysine residues of protein L WT, indicating their relative solvent exposure and potential accessibility.

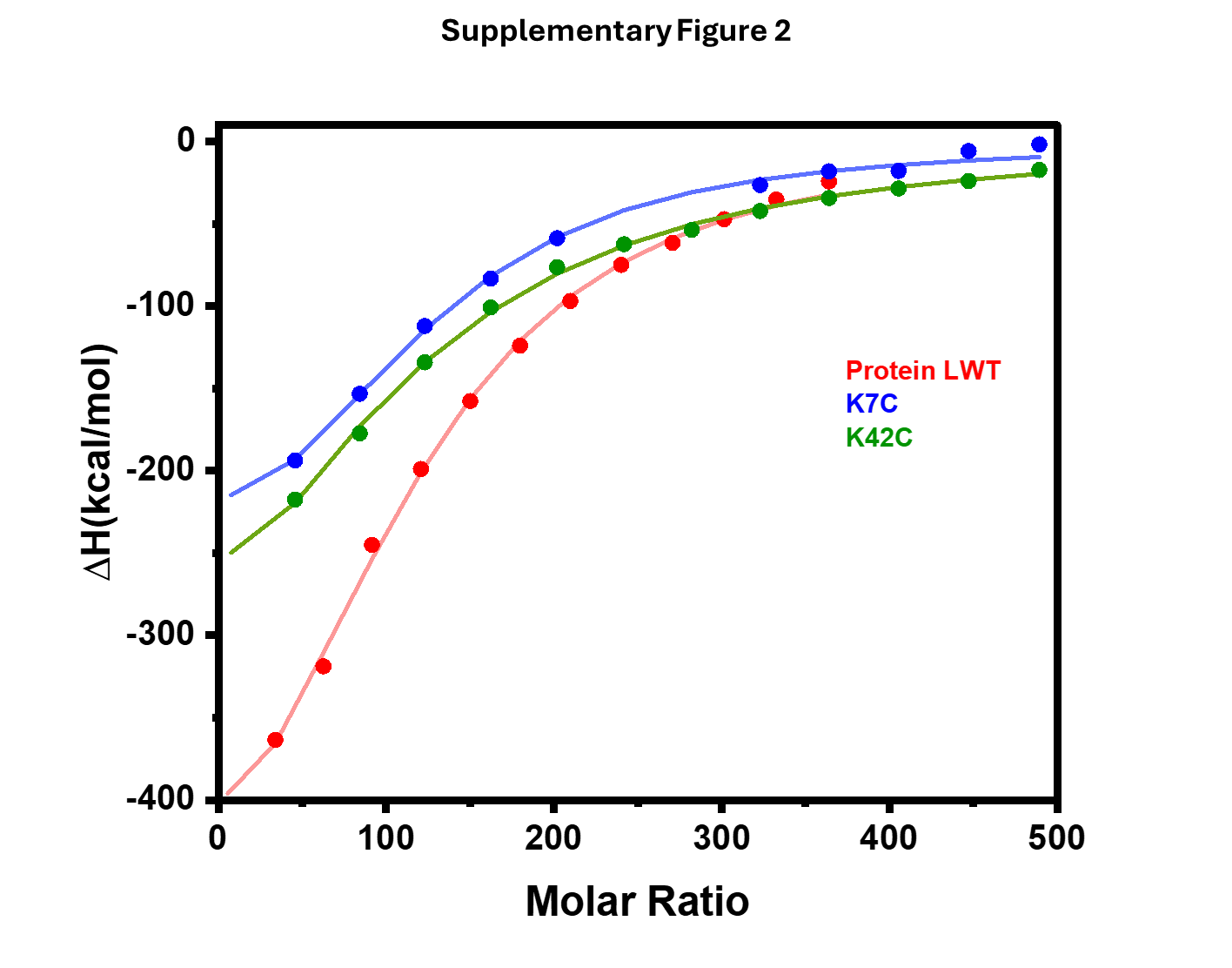

**SI Fig. 1.** Isothermal titration calorimetry (ITC) binding isotherms showing the enthalpy change (ΔH) as a function of molar ratio for the interaction of GNPs with wild-type protein L WT (red), and cysteine mutants K7C (blue) and K42C (green).

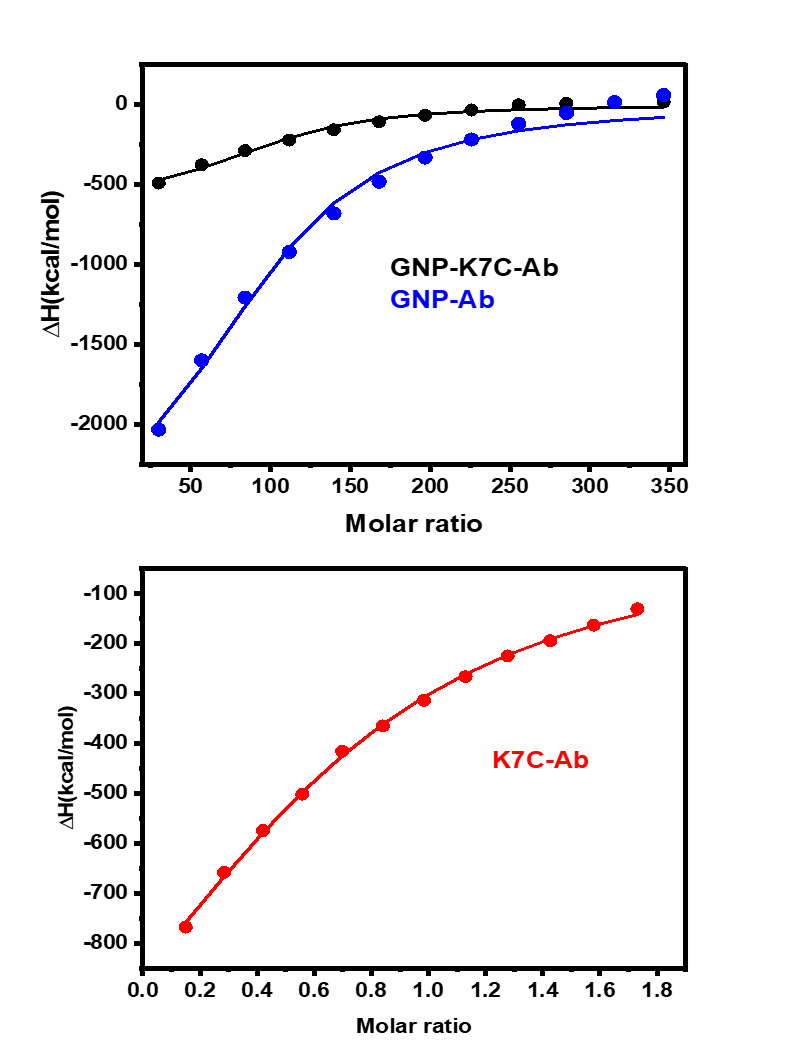
**SI Fig. 2.** Binding isotherms, plotting the heat change per mole of injectant (Antibody) (kcal/mol) against the molar ratio of GNP (blue), K7C (red) and GNP coated with K7C (black).

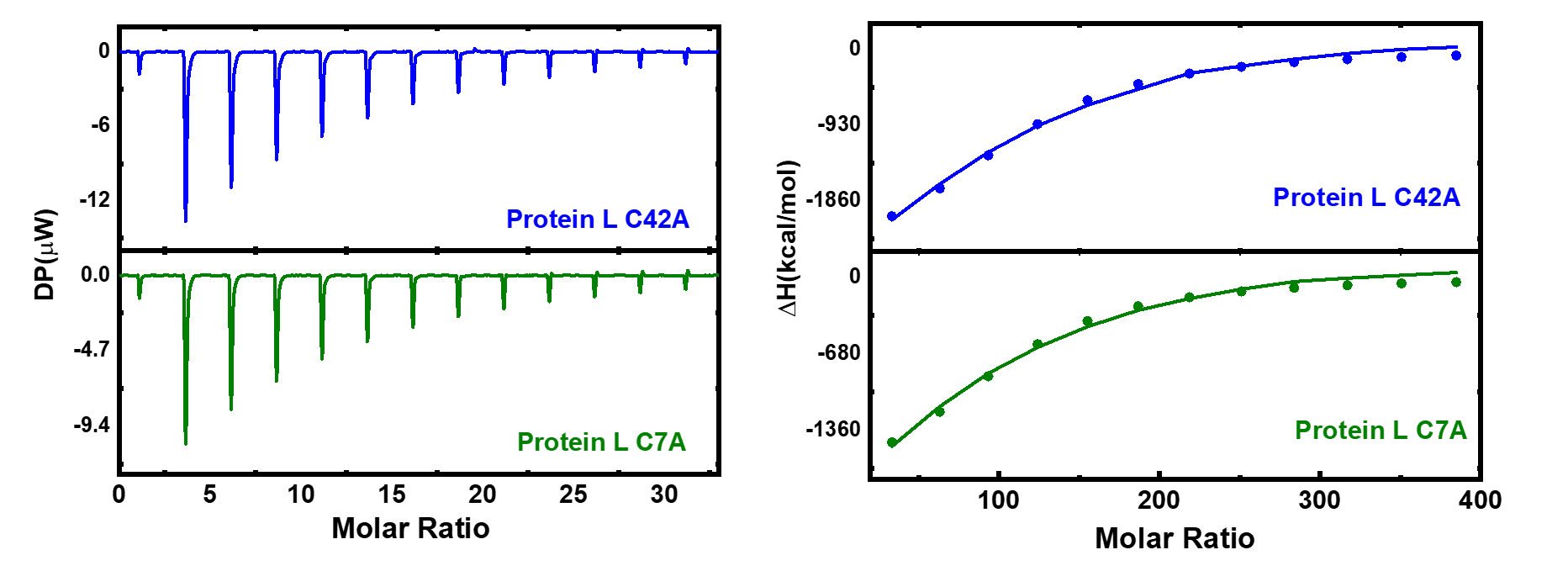

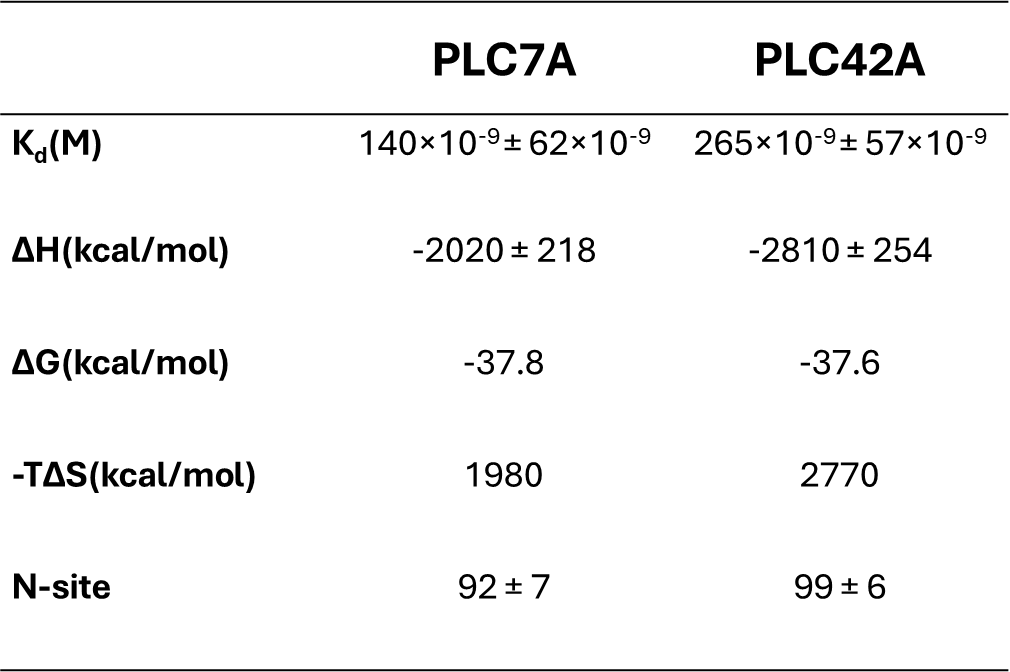
**SI Fig. 3.** ITC experiments probing the binding energetics of protein L mutants (K7/42A) onto GNPs. Corrected differential power (DP) vs. time plot of protein L K42A (blue) and K7A (green), titrated against GNPs. The thermodynamic saturation time points for the titrations are similar, with nearly identical exothermic responses, as indicated by the DP values (right panel). Binding isotherms, plotting the heat change per mole of injectant protein K42A (blue) and K7A (green) (kcal/mol) against the molar ratio of GNP.

**SI Table 2.** Thermodynamic parameters of protein L K7/42A interaction with GNPs, as obtained from the ITC titration.

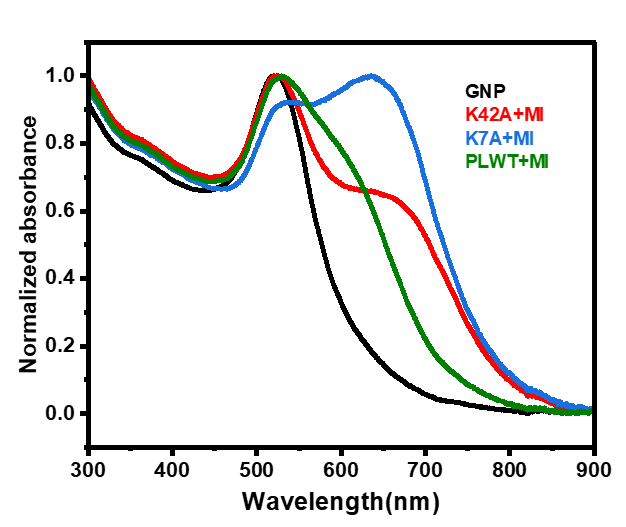
**SI Fig. 4.** UV-Visible absorption spectra upon addition of 2-MI to protein L K7A/K42A, and WT incubated with GNPs for 24 hr.

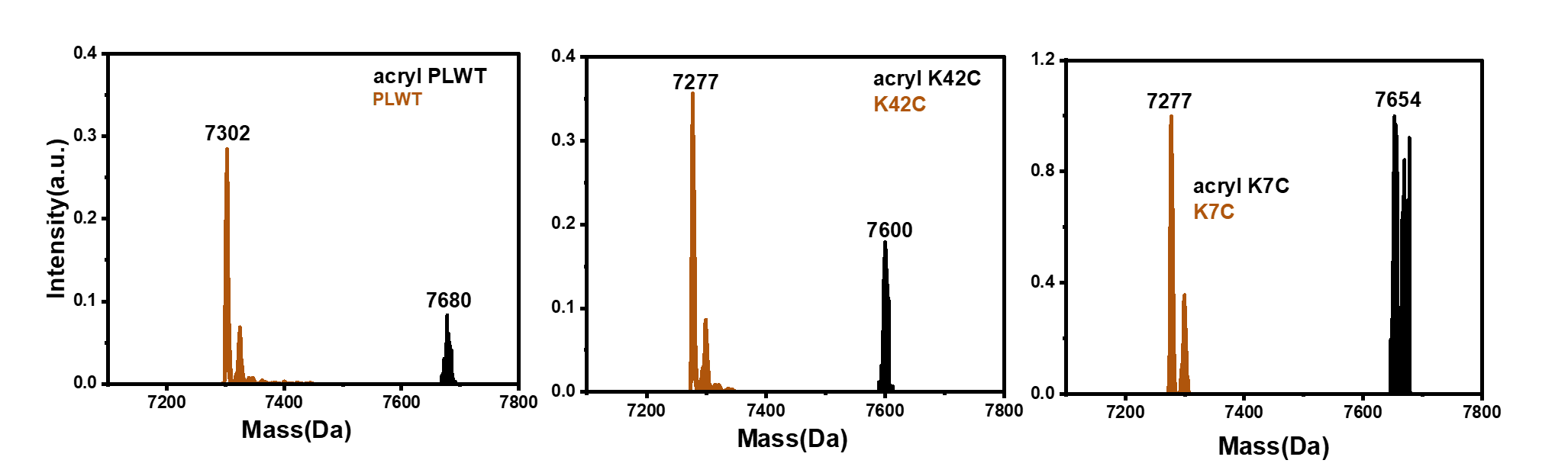
**SI.1 ESI–MS analysis of Protein L variants after acrylation**

**SI Fig. 5.** (A) PLWT, (B) K42C, and (C) K7C ESI-MS showing distinct mass shifts corresponding to acrylation of lysine residues. Acrylation was carried out using NHS-acrylate, resulting in approximately five, four, and five acrylates in PLWT, K7C, and K42C, respectively.

We evaluated the role of cysteine in protein L–GNPs binding by blocking lysines through NHS-ester acrylation. We used molar masses: NHS acrylate 169 g/mol, acrylate 71 g/mol, and protein L 7.114 kDa; these guided estimates of the maximum acrylations per protein. ESI-MS was used to quantify modifications: PLWT carried five acrylations, K42C four, and K7C five. Because protein L has seven lysines, the ~two unmodified sites are likely buried or sterically restricted, and thus inaccessible to the reagent. Their poor reactivity suggests minimal participation in Protein L–GNP interactions. Selective lysine blocking therefore isolates cysteine-mediated binding and maps which lysines are surface accessible under our conditions. Overall, this study indicates that accessible lysines dominate the initial contact with the negatively charged GNP surface. Cysteine-dependent effects become evident only after these lysines are masked. This framework enables clearer attribution of binding roles across variants and supports subsequent mechanistic comparisons.

**SI.2 Reversibility of acrylated Protein L variants incubated with GNPs**

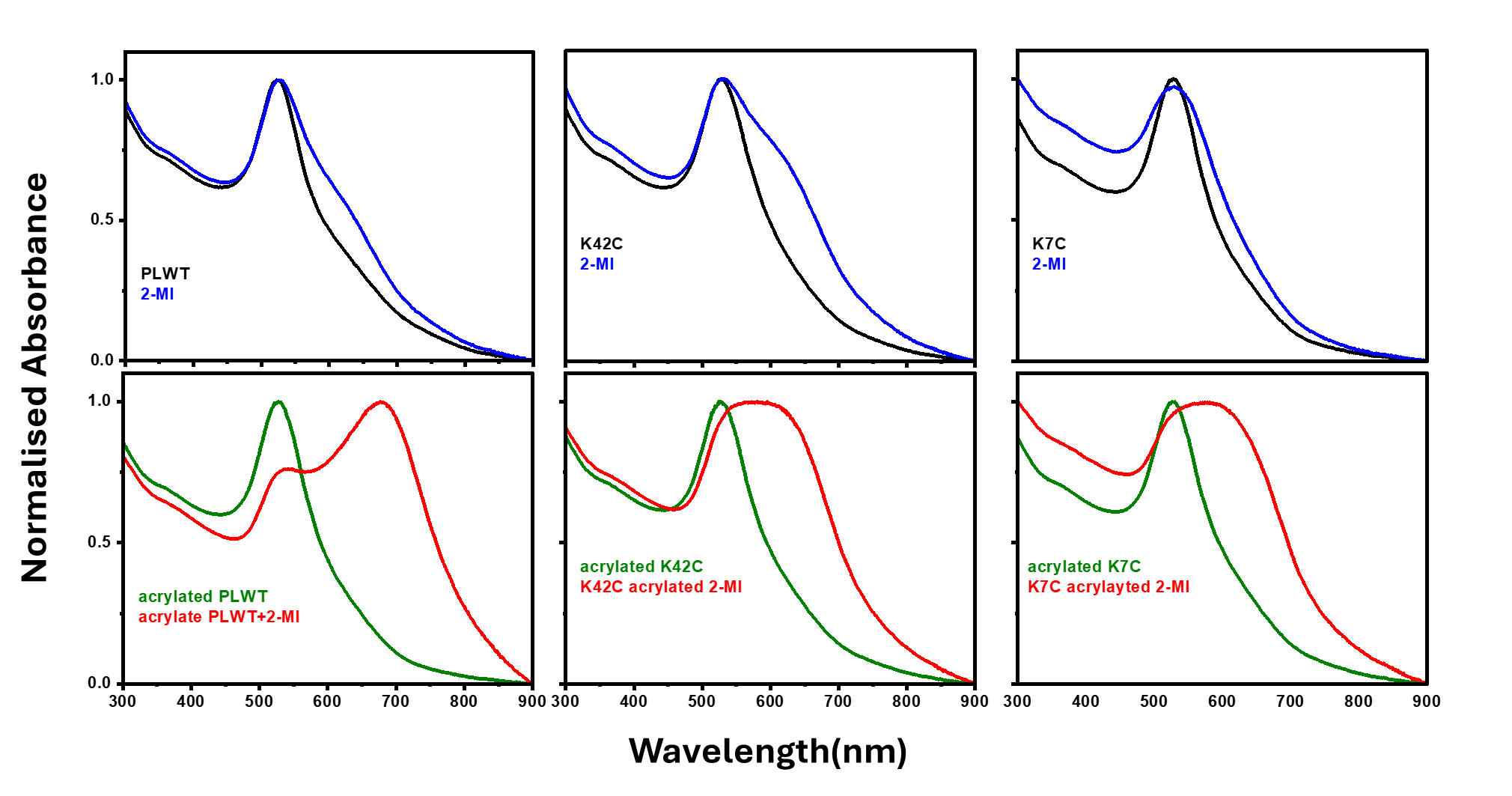
**SI Fig. 6.** UV–Vis absorption spectra of protein L variants with GNPs and 2-MI. (A) PLWT, (B) K42C, and (C) K7C were incubated with GNPs for 24 h. Black and blue curves correspond to non-acrylated proteins in the absence and presence of 2-MI, respectively. Red and green curves represent acrylated proteins in the absence and presence of 2-MI, respectively. Acrylation blocks lysine residues, allowing specific evaluation of cysteine–GNP interactions. Acrylation increases aggregation of PLWT, indicating that lysines contribute significantly to GNP capping; K7C and K42C show similar aggregation after acrylation, highlighting cysteine contribution in K42C, after lysine is acrylated.

Acrylation was used to block lysine residues so that their role in binding to GNPs was minimized, allowing us to focus on cysteine-mediated effects. Binding reversibility was tested with 2-MI. If capping is reversible, 2-MI promotes aggregation, seen as broadening of the SPR peak near ~520 nm. PLWT has no cysteine. After 24 h with GNPs, it showed very little aggregation, indicating stable capping driven by lysines. When PLWT was acrylated, strong aggregation and marked SPR broadening was seen, confirming that lysines are the major contributors to PLWT-mediated capping.

K7C contains an exposed lysines and a surface-exposed cysteine. In its non-acrylated form, it immobilised on GNPs and showed minimal aggregation, suggesting cooperative interaction via lysines and cysteine. In contrast, K42C has an exposed cysteine but lacks an exposed lysine to initiate contact with the negatively charged GNP surface. It therefore aggregated substantially even without acrylation, indicating weaker stabilisation. After acrylation, both K7C and K42C showed aggregation. Notably, their acrylated forms behaved similarly, each showing SPR broadening around 520 nm. This aggregation was less than that of acrylated PLWT yet nearly identical between K7C and K42C, indicating that once lysines are blocked, both variants rely mainly on cysteine for interaction and display comparable capping efficiency.
